# HDLs extract lipophilic drugs from cells

**DOI:** 10.1101/2020.12.08.415984

**Authors:** Adi Zheng, Gilles Dubuis, Carla Susana Mendes Ferreira, Thomas Mercier, Laurent Decosterd, Christian Widmann

## Abstract

High-density lipoproteins (HDLs) prevent cell death induced by a variety of cytotoxic drugs. The underlying mechanisms are however still poorly understood. Here we present evidence that HDLs efficiently protect cells against thapsigargin (a SERCA inhibitor) by extracting the drug from cells. Drug efflux could also be triggered to some extent by low-density lipoproteins (LDLs) and serum, which contains lipoproteins. HDLs did not reverse the non-lethal mild endoplasmic reticulum (ER) stress response induced by low thapsigargin concentrations or by SERCA knock-down but HDLs inhibited the toxic SERCA-independent effects mediated by high thapsigargin concentrations. HDLs were also found to extract other lipophilic compounds, such as the anti-diabetic drug glibenclamide. In contrast, hydrophilic substances (doxorubicin hydrochloride, rhodamine 123) were not extracted from cells by HDLs. This work shows that HDLs utilize their capacity of loading themselves with lipophilic compounds, akin to their ability to extract cellular cholesterol, to reduce the cell content of hydrophobic drugs. Silencing of the P-glycoprotein/ABCB1 transporter reduced the capacity of cells to load thapsigargin on HDLs. This work suggests that HDL-mediated cell efflux of toxic lipophilic xenobiotic is beneficial but also that HDL-mediated efflux can be detrimental to the therapeutic benefit of lipophilic drugs such as glibenclamide. Lipoprotein-mediated drug efflux should therefore be considered when evaluating drug efficacy.

## Introduction

HDLs possess multifaceted biological protective properties including anti-inflammation, anti-oxidation and anti-apoptosis, which can be beneficial for the treatment of diseases [1]. The mechanism(s) used by HDLs to protect cells are poorly understood [2]. Sphingosine-1-phosphate receptor-mediated Akt signaling has been proposed to be a pathway triggered by HDLs that protects cells [3]. However, a recent study [4] found no evidence for Akt being involved in HDL-mediated cell protection. How HDLs protect cells remains therefore largely unexplained.

HDLs inhibit endoplasmic reticulum (ER) stressor-induced death [2]. They do so via different mechanisms depending on which ER stressors are used. For example, in beta cells, HDLs blocked the ability of the sarco/endoplasmic reticulum Ca^2+^-ATPase (SERCA) pump inhibitor thapsigargin (TG) from inducing a strong unfolded protein response (UPR) and subsequently cell death [5] but HDLs did not prevent the inhibitory action of TG on SERCA [5]. On the other hand, while HDLs protected beta cells from the protein glycosylation inhibitor tunicamycin, it did so with minimal impact on the UPR[6]. Hence, in one case HDL-mediated protection was associated with UPR inhibition while in the other, protection occurred despite activation of the UPR. This indicates that HDLs can activate more than one anti-death pathway in cells.

A well-known property of HDLs is their capacity to extract cholesterol from cells. HDLs can also bind to drugs, especially hydrophobic molecules [7], and act as drug carriers [8, 9]. Whether the binding capacity of HDLs to particular drugs translates into an ability to extract the drugs from cells has not been tested yet. This prompted us to investigate whether HDLs can extract drugs from cells and whether this protects cells from drugs with cytotoxic properties.

## Materials and methods

### Reagents and Antibodies

SERCA2 antibody (ref 9271s; lot 14) was obtained from Cell signaling. Alexa Fluor 680 goat anti-rabbit antibody (ref A21109, lot 1816534) and Lipofectamine RNAiMAX transfection reagent (ref 13778030) were from ThermoFisher. BODIPY-FL thapsigargin and BODIPY-FL Glibenclamide were purchased from Marker Gene and Life Technology (Ref M4700; Lot 291AAN029, Ref E34251; Lot 2069641 respectively). Hoechst 33342 is from Thermofisher (Ref H3570). Doxorubicin hydrochloride was from Sigma-Aldrich (Ref: 44583-1MG), Rhodamine 123 was purchased from Sigma-Aldrich (Ref: 83702-10MG). The siPOOLs directed at ABCA1(lot:19-1-001), ABCB1 (lot:5243-1-001), ABCG1 (lot:9619-1-001) and ABCG2 (lot:9429-1-001), and a non-specific control siPOOL (lot:N000-051), were purchased from Biotech. The siPOOLs are high complexity pools of 30 optimally-designed siRNAs [10].

### Lipoprotein isolation

HDLs and LDLs were prepared from human serum by sequential density ultracentrifugation [11, 12].

### Cells and cell culture

Wild-type DLD1 (gift from Prof. Bert Volgenstein at the Core Cell center Baltimore) cells and Hela cells were maintained in RPMI 1640 (Gibco; ref 61870-010; lot 1880320) supplemented with 10% FBS (Gibco; ref 10270-106; lot 42G5062K) at a temperature of 37°C with 5% CO_2_. HEK293T cells, MCF7 cells and mouse embryonic fibroblasts (MEFs) were maintained in DMEM (Gibco; ref 61965-059;) supplemented with 10% FBS (Gibco; ref 10270-106; lot 42G5062K) at a temperature of 37°C with 5% CO_2_. MIN6 clone B1 mouse insulinoma cells (kindly provided by Dr. P. Halban, University Medical Center, Geneva, Switzerland) were cultured in high-glucose DMEM (Gibco; ref 61965-026) supplemented with 15% FBS, 1 mM of sodium pyruvate (Gibco; ref 11360-070) and 70 μM freshly added beta-mercaptoethanol (Gibco; ref 31350-010) at a temperature of 37°C with 5% CO_2_.

### Extracellular and intracellular drug quantitation

DLD-1 cells were plated in 6 well plates at a density of 500’000 cells per well and cultured for 24 hours in 2 ml of culture medium. The cells were then treated with 20 μM thapsigargin or 100 nM staurosporine in 1 ml fresh media for 2 hours. Cells were washed with PBS thrice and then incubated in fresh RPMI, 10% FBS in the absence (control) or in the presence of 1 mM HDL. Two hours later, media were collected in Eppendorf tubes. Cells were washed with PBS and then trypsinized. Cell pellets were kept in Eppendorf tubes. The drug content in the samples was analyzed by high performance liquid chromatography coupled to tandem mass spectrometry (HPLC-MS/MS).

### Drug content analysis by mass spectrometry

#### Incubation media sample preparation

a 300 μl aliquot of the incubation medium sample was mixed with 300 μl MeOH. This mixture was then vortex-mixed, and centrifuged at 16’000g for 10 minutes. The supernatant (500 μl) was transferred into an HPLC glass vial.

#### Cell sample preparation

cells were mixed with 300 μl MeOH. This cellular suspension was sonicated for 30 minutes. H_2_O (300 μl) was then added, and the sample centrifuged at 16’000 g for 10 minutes to eliminate solid cellular debris. The supernatant (500 μl) was transferred into an HPLC glass vial.

#### Calibration curves

quantitative analysis of the concentrations was performed using the external standard method. A calibration standard curve was calculated and fitted by quadratic log-log regression of the peak areas. The lower limit of quantification were 50 ng/mL for thapsigargin and 1 ng/mL for staurosporine.

### Intracellular drug quantitation by flow cytometry

Cells were plated in 6 well plates at a density of 150’000 cells per well in 2 ml RPMI, 10% FBS and cultured for 24 hours. Then, the medium was replaced with 1 ml fresh medium containing or not the indicated BODIPY-labelled drugs or naturally fluorescent drugs for 1 hour. The cells were then washed once and then left untreated or incubated with the indicated lipoproteins or proteins for various periods of time. Finally, cells were washed once with 1 ml PBS and trypsinized with 120 μl of trypsin/EDTA for about 2 minutes and recovered following the addition of 500 μl of RPMI, 10% FBS, centrifugated at about 200 g for 3 minutes and finally resuspended in 500 μl PBS for flow cytometry analysis. The fluorescence of BODIPY-TG, BODIPY-glibenclamide, doxorubicin hydrochloride and Rhodamine 123 fluorescence was measured using the KO525, FITC, PE, and FITC channels of a CytoFlex-S flow cytomter (Beckman), respectively. Data analysis was done with Kaluza Version 1.3 software.

### siRNA transfection

The first round of siPOOL transfection was performed at the time of cell seeding (80,000 cells/well, 400 μl DMEM, 10% FBS) in 12-well plates. The transfection mix was made as follows: 50 μl Opti-MEM (Thermo 11058021) were placed in two different sterile Eppendorf tubes, 1.5 μl siRNA (5 μM stock) were added to one tube and 1.25 μl RNAi-MAX to the other. Then the content of the two tubes were mixed together and incubated at room temperature for 5 minutes. The mixture was added dropwise in the well of the plates. Twenty-four hours later, the media was replaced with 0.4 ml fresh media and another siRNA transfection was performed. Cells were analyzed 48 hours or 72 hours after the first transfection as indicated.

### HEK293T cell number counting

Following siPOOL transfection, HEK293T cells were treated with 20 μM TG for 24 hours in the absence or in the presence of 1 mM HDLs. The dead floating cells were removed carefully and the attached cells were collected in 1 ml 10% DMEM medium and centrifuged at about 200 g for 3 minutes. The cell pellets were then suspended in 0.5-1 ml of DMEM, 10% FBS. A volume of 10 μl was taken and placed on a hemocytometer and cells were counted. The total cell number for each condition were then calculated based on the suspension medium volume.

### Real-time PCR

Cells were treated with unlabeled drugs in an otherwise similar manner as described in the previous section except that the pelleted cells after trypsinization and centrifugation were frozen at −80°C until processed for RNA extraction. Total RNA was extracted from cells using High pure RNA isolation kit from Roche (Ref. 11828665001, lot. 38800800) according to the manufacturer’s instructions. Reverse transcription from RNA to cDNA was performed with the Transcriptor Universal cDNA Master kit from Roche (Ref. 05893151001, lot. 32966400). The semiquantitative real-time PCR was proceeded with the faststart universal SYBR green master from Roche (Ref. 04913914001, lot. 11929100) using gene-specific primers. Data were normalized to mRNA of GAPDH as a housekeeping gene and were analyzed by the 2^−ΔΔCt^method. Relative expression of genes was expressed as fold change over control. The following primers were used for the RT-PCR analysis: h-SERCA2 F (#1612): ATG GGG CTC CAA CGA GTT AC (nucleotides 648-667 of human SERCA2, variant a; NCBI entry NM_001681.4; identical sequence in human variant b); h-SERCA2 R (#1613): TTT CCT GCC ATA CAC CCA CAA (nucleotides 851-871 of human SERCA2, variant a; NCBI entry NM_001681.4), h-BIP F (#1617): GAA AGA AGG TTA CCC ATG CAG T (nucleotides 702-723 of human BIP; NCBI entry NM_005347.5); h-BIP R (#1618): CAG GCC ATA AGC AAT AGC AGC (nucleotides 830-850 of human BIP; NCBI entry NM_005347.5), h-XBP1s F (#1619): TGC TGA GTC CGC AGC AGG TG (nucleotides 528-547 of human XBP1,variant 2; NCBI entry NM_001079539.2); h-XBP1s R (#1620): GCT GGC AGG CTC TGG GGA AG (nucleotides 677-696 of human XBP1,variant 2; NCBI entry NM_001079539.2) [13], h-CHOP F (#1621): GGA AAC AGA GTG GTC ATT CCC (nucleotides 491-511 of human CHOP, variant 5; NCBI entry NM_004083.6; identical sequence in variants 1-4 and 6); h-CHOP R (#1622): CTG CTT GAG CCG TTC ATT CTC (nucleotides 586-606 of human CHOP, variant 5; NCBI entry NM_004083.6; identical sequence in variants 1-4 and 6). h-ABCB1 F (#1602): TTG CTG CTT ACA TTC AGG TTT CA (nucleotides 897-919 of human ABCB1,variant 1; NCBI entry NM_001348945.2; identical sequence in variants 2-4); h-ABCB1 R (#1603): AGC CTA TCT CCT GTC GCA TTA (nucleotides 981-1001 of human ABCB1,variant 1; NCBI entry NM_001348945.2; identical sequence in variants 2-4), h-GAPDH F (#1578): CTG ACT TCA ACA GCG ACA CC (nucleotides 873-892 of human GAPDH,variant 7; NCBI entry NM_001357943.2; identical sequence in variants 1-4); h-GAPDH R (#1579): TGC TGT AGC CAA ATT CGT TG (nucleotides 967-986 of human GAPDH, variant 7; NCBI entry NM_001357943.2; identical sequence in variants 1-4). h-ABCG2 F (#1606): CAG GTG GAG GCA AAT CTT CGT (nucleotides 350-370 of human ABCG2, variant 7; NCBI entry NM_001348989.2; identical sequence in variants 1-6); h-ABCG2 R (#1607): ACC CTG TTA ATC CGT TCG TTT T (nucleotides 575-596 of human ABCG2, variant 7; NCBI entry NM_001348989.2; identical sequence in variants 1-6).

### Western blotting

Western blotting was performed as described previously [14]. Briefly, cells were lysed in mono Q-c lysis buffer. Protein concentrations in the cell lysates were quantified by Bradford assay. Equal amounts of protein per sample (20 μg) were loaded in a 10% polyacrylamide gel; proteins were then transferred to nitrocellulose membranes. Following Ponceau (0.1% Ponceau, 50% acetic acid) staining to confirm proper protein transfer, the membrane was blocked using 5% BSA in TBST for 30 minutes. The membrane was subsequently incubated with primary antibodies against SERCA2 (1:1000 in 5% BSA) overnight at 4°C, washed for 30 minutes in TBS-tween 20%, incubated with a secondary antibody (diluted 1:10000 in 5% BSA) for 1 hour at 5°C and washed for 30 minutes in TBS-Tween 20%. The blots were finally detected by Odyssey infrared imaging system (LICOR Biosciences, Bad Homburg, Germany)

### SERCA2 knockdown

The mSERCA2-shRNA2.pll3.7 vector (plasmid #748) was used to generate lentiviruses encoding an shRNA against SERCA2 (the sequence targeted by this shRNA is conserved between humans and mice). It was constructed by subcloning annealed oligonucleotides #912 and #913 into the pLentiLox3.7 lentiviral vector (plasmid #627) (Addgene cat. n°11795) opened with HpaI and XhoI. The sequences of the oligonucleotides were:

#912 (sense): 5’-T (used to complete the U6 promoter) GCAACTGTCTATTTCTGCT (nucleotides 3735-3753 of mouse SERCA2; NM_027838) TTCAAGAGA (shRNA loop) AGCAGAAATAGACAGTTGC (nucleotides 3753-3735 of mouse SERCA2; NM_027838) TTTTTT (linker) C (N1 nucleotide of the XhoI site)-3’.

#913 (anti-sense): 5’-TCGAG (N1-N5 nucleotides of the XhoI site) AAAAAA (linker) GCAACTGTCTATTTCTGCT (nucleotides 3735-3753 of mouse SERCA2; NM_027838) TTCTCTTGAA (shRNA loop) AGCAGAAATAGACAGTTGC (nucleotides 3753-3735 of mouse SERCA2; NM_027838) A (used to complete the U6 promoter)-3μ.

Recombinant lentiviruses were produced in HEK293T cells [15]. Briefly, cells were co-transfected using the calcium phosphate DNA precipitation method with the lentiviral mSERCA2-shRNA2.pll3.7 vector (plasmid #748), the envelope protein-coding plasmid (pMD2.G, plasmid #554) and the packaging construct (pSPAX2, plasmid #842). After a 6 hour transfection period, medium was replaced with fresh medium. Forty-eight hours later, the virus-containing medium was harvested and kept at −80 °C.

DLD-1 cells (20’000 cells per well) were seeded in 6-well plates. The following morning, cells were infected with lentiviruses encoding the shRNA directed against SERCA2. The infection rate was quantitated by monitoring GFP fluorescence after three days. The minimal volume of viruses leading to GFP expression in 100% of the cells (assessed by flow cytometry) was used in subsequent experiments.

### Cell viability (propidium iodide assay) in DLD-1 cells

After the indicated treatment, cells (including the floating cells in the medium) were collected and then incubated with 8 μg/ml propidium iodide (PI) in PBS for 5 minutes at room temperature. PI incorporation into cells was then analyzed by flow cytometry.

### Pycnosis assessment in Min6 cells

After the indicated treatment, Min6 cells were fixed with 4% paraformaldehyde (PFA) and incubated five minutes with 10 μg/ml Hoechst 33342. Pycnotic cells was then determined visually using fluorescence microscopy. At least 700 cells were counted for each condition.

### Data presentation and statistics

Data from independent experiments are depicted with symbols of different motifs and shading. Comparisons between multiple groups were performed using two-way ANOVA or one-way ANOVA followed by Dunnett’s multiple comparisons test using GraphPad Prism. Comparisons between two groups were performed using Student’s t-test in GraphPad Prism.

## Results

### The effect of HDLs on different TG concentrations

TG is a specific and irreversible inhibitor of the sarco/endoplasmic reticulum Ca^2+^-ATPase (SERCA) pump [16, 17]. SERCA deficiency induces ER dysfunction [18]. As shown previously [14], HDLs were able to protect cells from high concentration (≥10 μM) TG-induced death (Figure 1), even when added after cells were pre-loaded with TG (Figure S1). TG induced the mRNA expression of ER stress-related markers (BIP, CHOP, and XBP1s) in a concentration-dependent manner (Figure 2). HDLs decreased the mRNA expression of these genes in cells stimulated with high TG concentrations (10 μM) to the levels detected when lower TG concentrations (≤5 μM) were used alone. However, HDLs did not alter BIP, CHOP and XBP1s mRNA expression induced by TG concentrations ≤5 μM.

**Figure 1.**
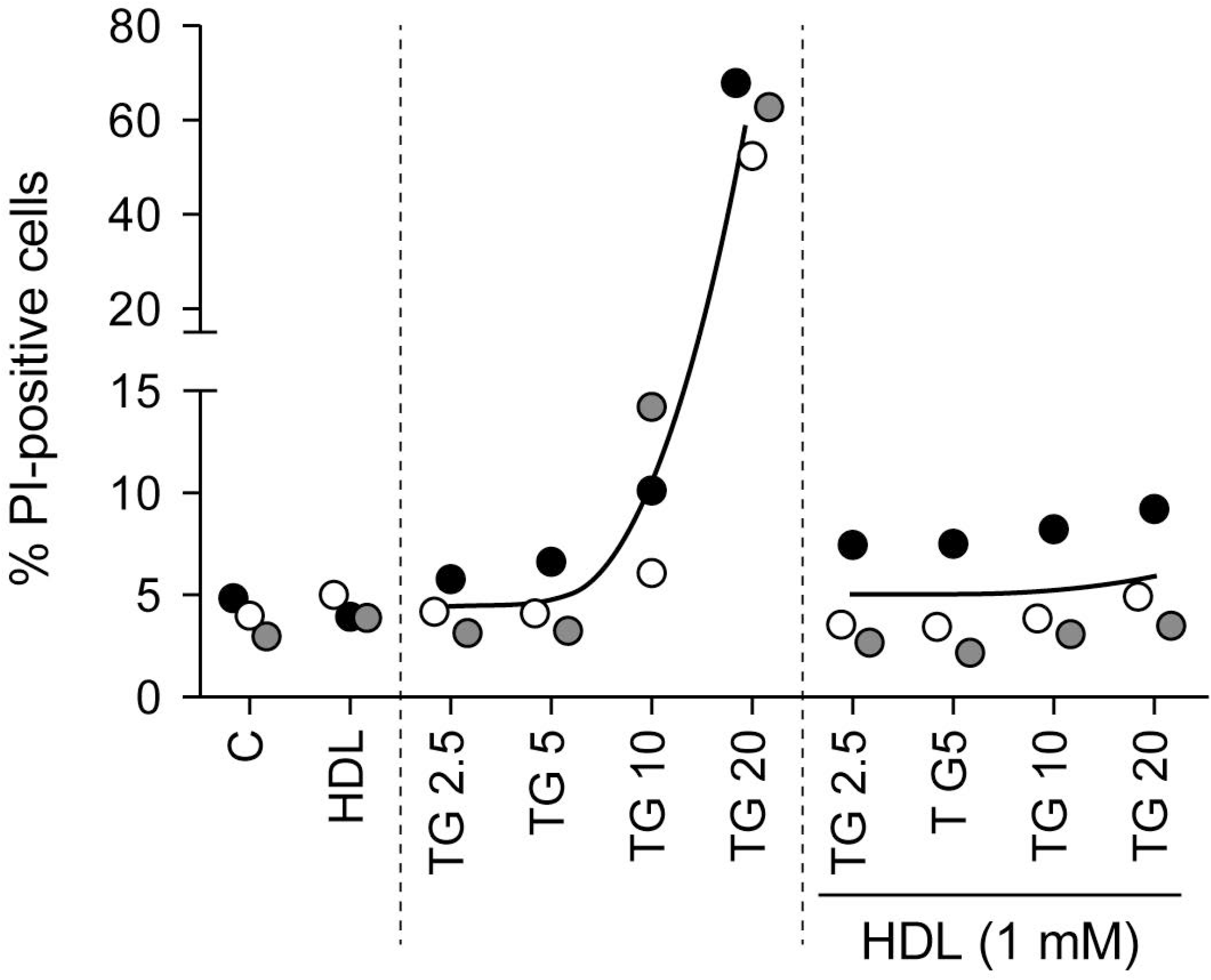
HDL-mediated inhibition of TG-induced cell death. DLD-1 cells were seeded in 6-well plates (100’000 per well). Twenty-four hours later, cells were treated for 48 hours with the indicated concentrations (in μM) of thapsigargin in the presence or in the absence of HDLs. Cell death was measured by flow cytometry following PI staining. Symbols with a given shade of grey are derived from a given independent experiment. The black curves go through the average values of the different experiments.

**Figure 2.**
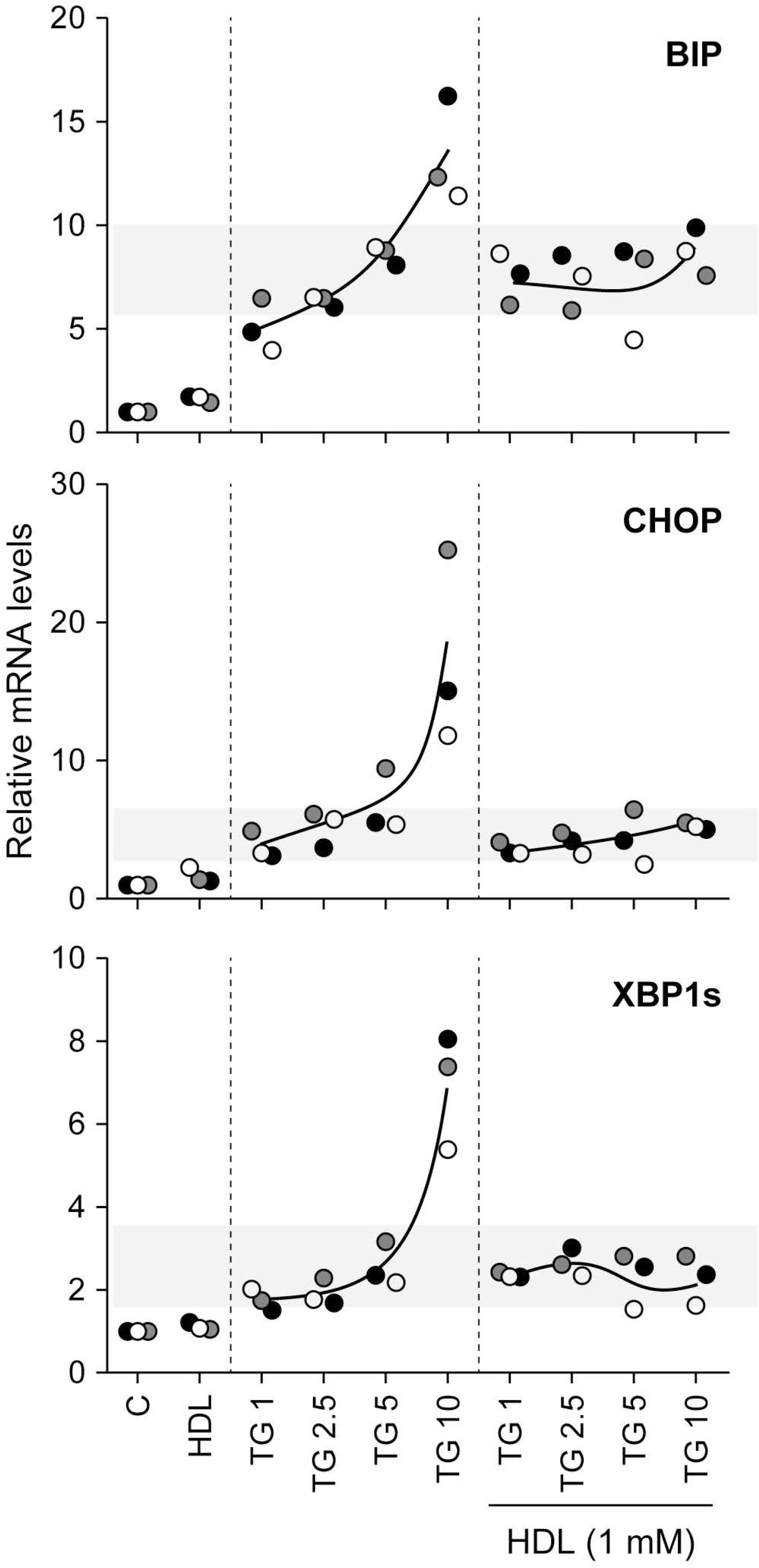
HDLs inhibit ER stress marker expression induced by high TG concentrations. DLD-1 cells were seeded in 6-well plates (200’000 per well). Twenty-four hours later, cells were treated for 24 hours with the indicated concentrations (in μM) of thapsigargin in the presence or in the absence of HDLs. The expression levels of UPR-induced proteins (BIP, CHOP, and XBP1s) were measured by RT-PCR.

### The effect of HDL on SERCA2 knockdown-induced ER stress

TG is a long-lasting irreversible inhibitor of SERCA but cells can adapt to this inhibition and restore ER homeostasis and protein translation and become refractory to TG effect [19]. The observation provided in Figure 2 indicates that HDLs do not alter the UPR response induced by concentrations of thapsigargin below 5 μM that are known to fully inactivate SERCA [17]. HDLs should therefore not affect the UPR response induced by SERCA invalidation. To assess this point, we knocked down SERCA2 in DLD-1 cells (SERCA1 and SERCA3 are not expressed in this cell line; Figure 3A). SERCA2 knockdown (Figure 3B) in DLD-1 induced mild ER stress, as indicated by the up-regulation of ER stress markers (Figure 3C) to levels equal or lower than those obtained with low concentrations of thapsigargin (compare Figure 2 and Figure 3). HDLs were not able to suppress the ER stress response induced by SERCA2 knockdown (Figure 3D). By themselves, HDLs slightly stimulated the expression of UPR genes, a likely consequence of their ability to modulate cholesterol levels in cells [20]. Despite their inability to prevent SERCA2 knockdown-induced mild ER stress response (Figure 3), HDLs were still capable of blocking the strong UPR response induced by high concentrations of TG in SERCA2 knockdown cells (Figure S2).

**Figure 3.**
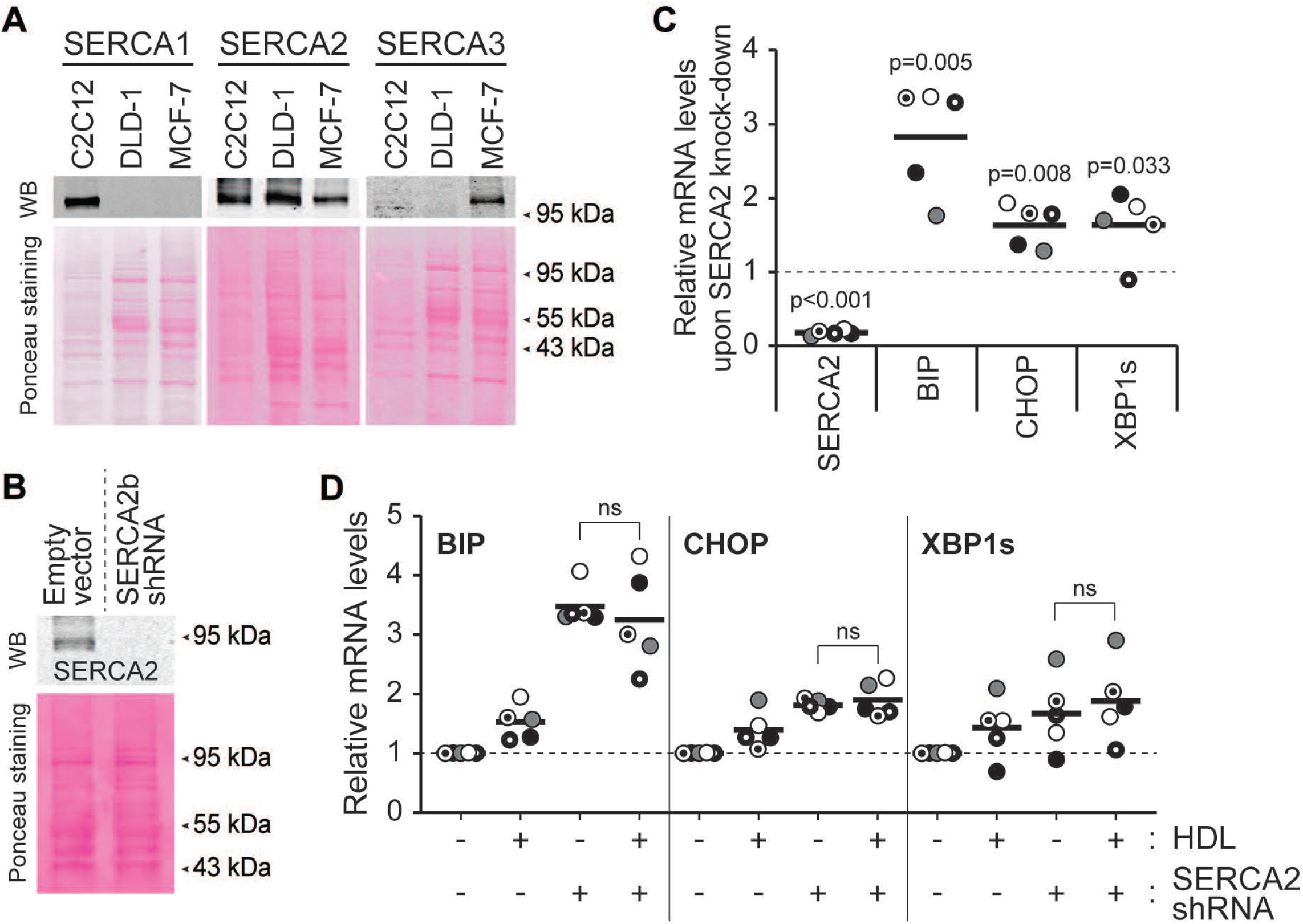
HDLs do not alleviate the stress response mediated by SERCA knock-down. **A**. SERCA1, SERCA2 and SERCA3 expression in DLD-1 cells was assessed by western blotting. C2C12 and MCF-7 cell lysates were used as positive controls for SERCA1 and SERCA3, respectively. **B**. Knockdown efficiency of SERCA2 in DLD-1 cells. **C**. The mRNA expression of SERCA2 and ER stress markers (BIP, CHOP and XBP1s) in SERCA2-knockdown cells. **D**. Effect of HDLs on the induction of ER stress marker mRNA expression induced by SERCA2 knockdown. In panels C and D, the results are normalized to the expression values obtained in untreated cells (dashed line). Symbols with a given shape and shading are derived from a given independent experiment. The black lines correspond to the mean.

We previously showed that HDLs could not prevent the transient increase of cytosolic calcium induced by the inhibitory action on SERCA [5]. Coupled with the present results, we can conclude that HDLs inhibit the SERCA-independent UPR and lethality induced by high concentrations of TG but that HDLs neither interfere with the ability of TG to inhibit SERCA nor with the subsequent ER mild stress response that ensues.

### HDLs extract lipophilic drugs from cells

A classical role of HDLs is to facilitate cholesterol efflux and to remove excess cholesterol from cells [1]. Conceivably, HDLs might have the capacity to decrease lipophilic drug content inside cells and therefore protect them against their lethal effects. To test this hypothesis, we determined whether HDLs could modulate the intracellular TG content. Figure 4A shows that HDLs decreased the amount of cell-associated TG and this was paralleled with an increase of TG in the HDL-containing medium. The ability of HDLs to extract TG from cells was also evidenced using a fluorescent version of TG (BODIPY-TG) (Figures 5 and S3 to S5). HDLs were able to extract other lipophilic drugs such as staurosporine (Figure 4B) and glibenclamide (Figure 6). In contrast to lipophilic drugs, hydrophilic drugs or compounds like doxorubicin hydrochloride, an anticancer drug, or Rhodamine 123, an ABCB1/p-glycoprotein activity sensor, were not extracted from cells by HDLs (Figure 6). Analyzing the physicochemical properties of the drugs used here (plus cholesterol as a classical HDL-interactor) using the SwissADME web tool [21] substantiated the notion that HDLs have the capacity to bind lipophilic but not hydrophilic molecules (Figure S6).

**Figure 4.**
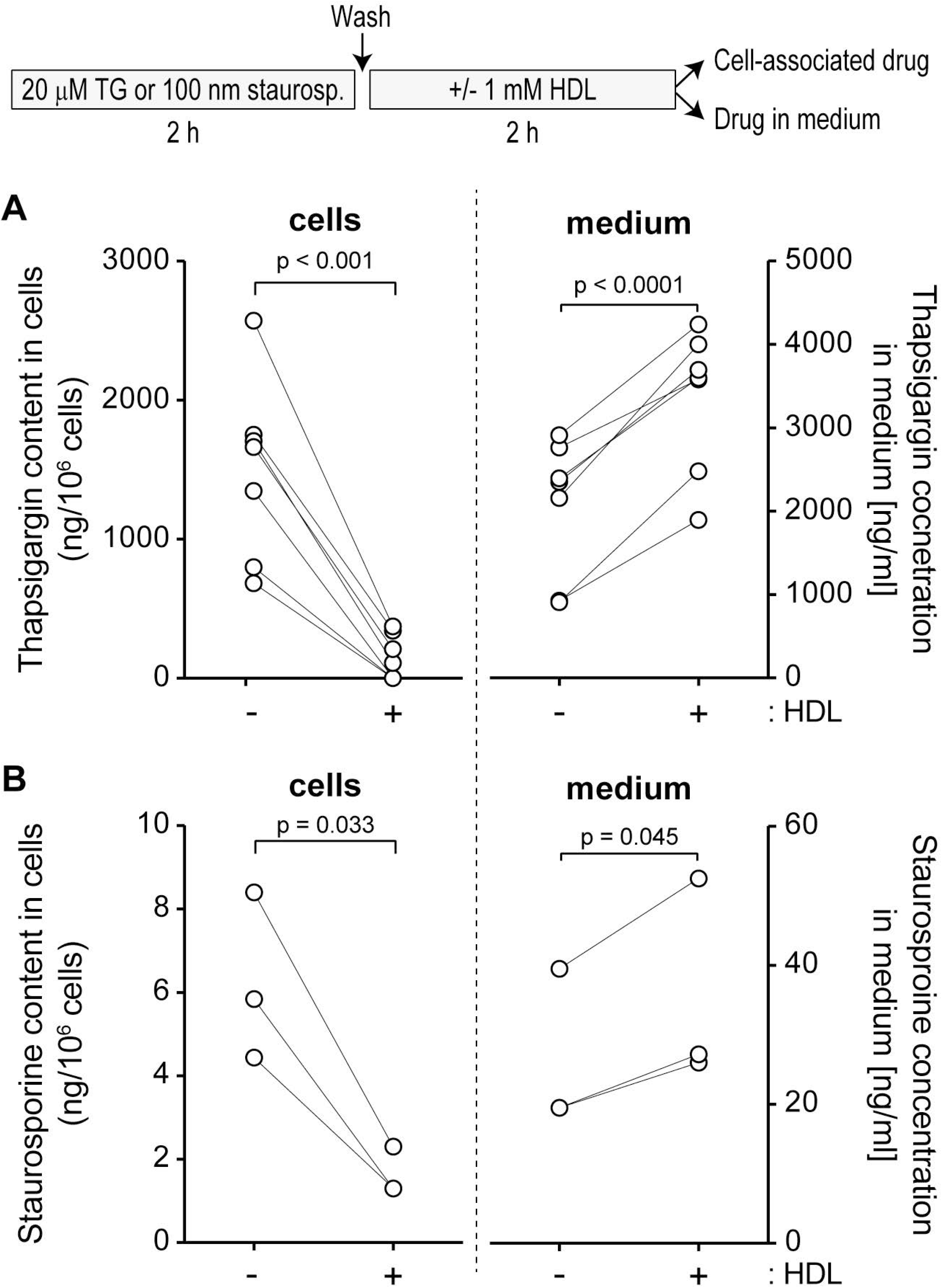
HDLs extract thapsigargin and staurosporine from cells. DLD1 cells (500’000 cells) plated in 6-well plates were treated 24 hours later as depicted in the scheme above the figure (see also the Methods). The cell-associated drug content and the drug found in the medium were then quantitated as described in the methods. Panels **A** and **B** describe the results obtained for thapsigargin and staurosporine, respectively.

**Figure 5.**
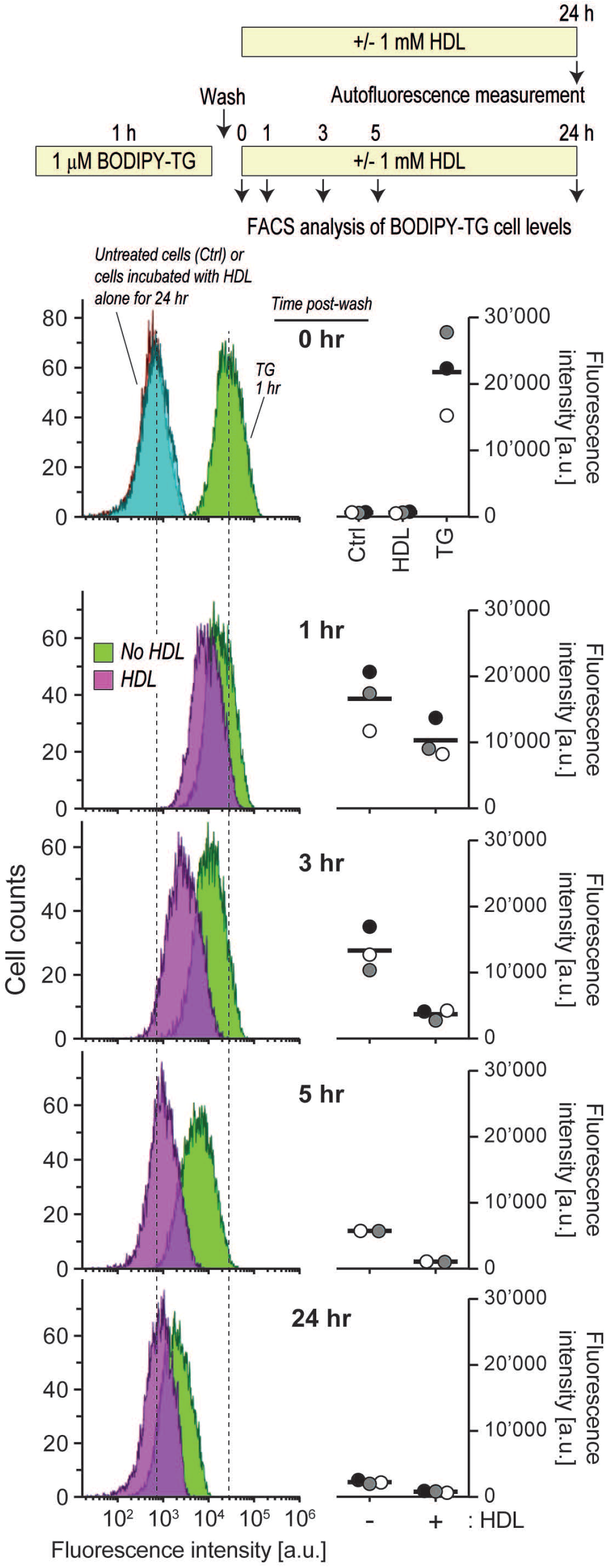
Kinetics of spontaneous and HDL-mediated thapsigargin cell release. Min6 cells (300’000 cells per well) were seeded in 6-well plates and cultured overnight. After treatment with 1 μM BODIPY-TG for 1 hour, cells were washed with PBS once and then incubated with medium or HDLs for the indicated periods of times before their drug content analyzed by flow cytometry. In the “0 hr” panel, the autofluorescence of untreated cells and cells incubated with HDLs alone for 24 hours are also shown (the two corresponding distributions fully overlap), as well as the profile of TG-BODIPY - incubated cells right after the washing step. The cells’ peak autofluorescence and the peak fluorescence intensity of 1 hour TG-BODIPY-incubated cells are indicated by dashed lines. The quantitation of 2-3 independent experiments (labelled with different symbols) are depicted on the right-hand side of the flow cytometry graphs. The latter are derived from the experiment corresponding to the grey symbols. The bars represent the mean of the cytometry distribution profiles.

**Figure 6.**
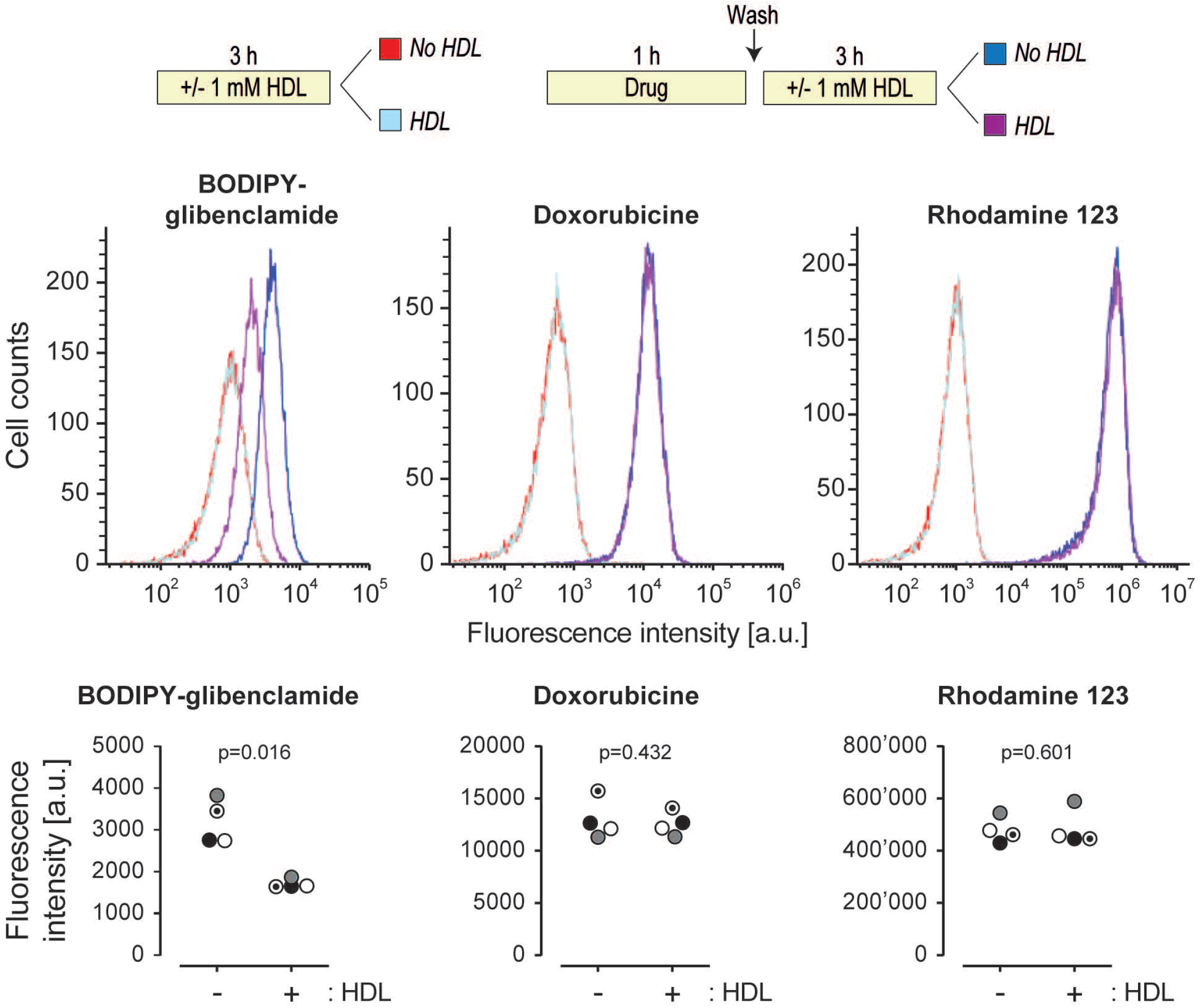
Effect of HDLs on BODIPY-glibenclamide, doxorubicin hydrochloride and rhodamine 123 efflux. Hela cells (120’000 per well) were plated in a 6-well plates. Then cells were treated as indicated in the scheme above the figures. Drug-associated fluorescence in cells was assessed by flow cytometry. The quantitation (mean of the cytometry distribution profiles) of 4 independent experiments (labelled with different symbols) are depicted below the flow cytometry profiles.

### Capacity of serum components other than HDLs to extract TG from cells

Lipoproteins, as well as proteins with the ability to bind lipophilic compounds such as albumin [22], are present in the serum used to supplement culture media. Conceivably, increasing the serum concentration in the media could negatively impact the ability of TG to accumulate in cells. This is indeed what was observed in two different cell lines (Figure S7A and S7B). Albumin alone was able to extract TG from cells but this accounted for only a fraction of what serum could achieve (Figure S7B). For example, there was ~50% less TG in cells incubated with 5% serum compared to cells incubated in the absence of serum but the decrease was only ~10% when the concentration of BSA found in 5% serum was used (20 μM; see tan region in Figure S7B). LDLs were also able to extract fluorescent TG from cells but again less efficiently than HDLs (Figure S7C). Moreover, HDLs were also more potent than LDLs to protect cells from the deleterious action of TG (Figure S7D). Altogether these results indicate that there are several serum components that can bind to lipophilic drugs such as TG but that none are as potent as HDLs to do so. The dose response curves obtained with serum or with the corresponding HDL concentrations are similar (compare the open circle curves in panels B and C of Figure S7). This indicates that HDLs explain most of the TG cell extracting activity of serum.

### ABCB1 participates in HDL-mediated drug efflux

To further elucidate the molecular mechanism of drugs export by HDLs, we examined the role of several transporters or receptors that have been related to cholesterol transport [23]. SR-BI is a surface protein HDLs can dock to [24] and that may therefore participate in drug efflux to these lipoproteins. We tested this hypothesis using SR-BI knock-out cells (Figure S8A). These cells have the same capacity as control cells to take up BODIPY-TG (Figure S8B). HDL-mediated BODIPY-TG efflux was not altered by the absence of SR-BI (Figure S8C), a finding consistent with the observation that SR-BI gene invalidation does not prevent HDLs from protecting cells against TG [5]. These results suggest that SR-BI plays no role in the ability of HDLs to extract lipophilic drugs from cells.

The ABCA1, ABCG1, ABCB1 and ABCG2 transporters can protect cells from the toxic effect of endogenous as well as xenobiotic molecules by mediating their efflux from cells [25–27]. We aimed to use siRNA-mediated silencing to test their involvement in HDL-mediated drug efflux. Unfortunately, the silencing efficiency for ABCA1 (~50% knockdown) and ABCG1 (~40% knockdown) was too weak, preventing us from testing their involvement in HDL-mediated drug efflux. On the other hand, ABCB1 and ABCG2 silencing was relatively efficient (Figure 7A-B and Figure 8D). ABCB1 knock-down partially inhibited the ability of the HDL to promote BODIPY-TG efflux (Figure 7C) and impaired the capacity of HDLs to protect cells from TG-induced death (Figure 7D). In the absence of HDLs, TG-induced death was also exacerbated by ABCB1 silencing, a likely consequence of lipoproteins in the serum being less able to extract the drug from cells (see Figure S7). In contrast to what was observed for ABCB1, ABCG2 silencing had no impact on the drug efflux capacity of HDLs (Figure S8D-E). These data suggest that ABCB1 mediates some of the drug efflux to HDLs.

**Figure 7.**
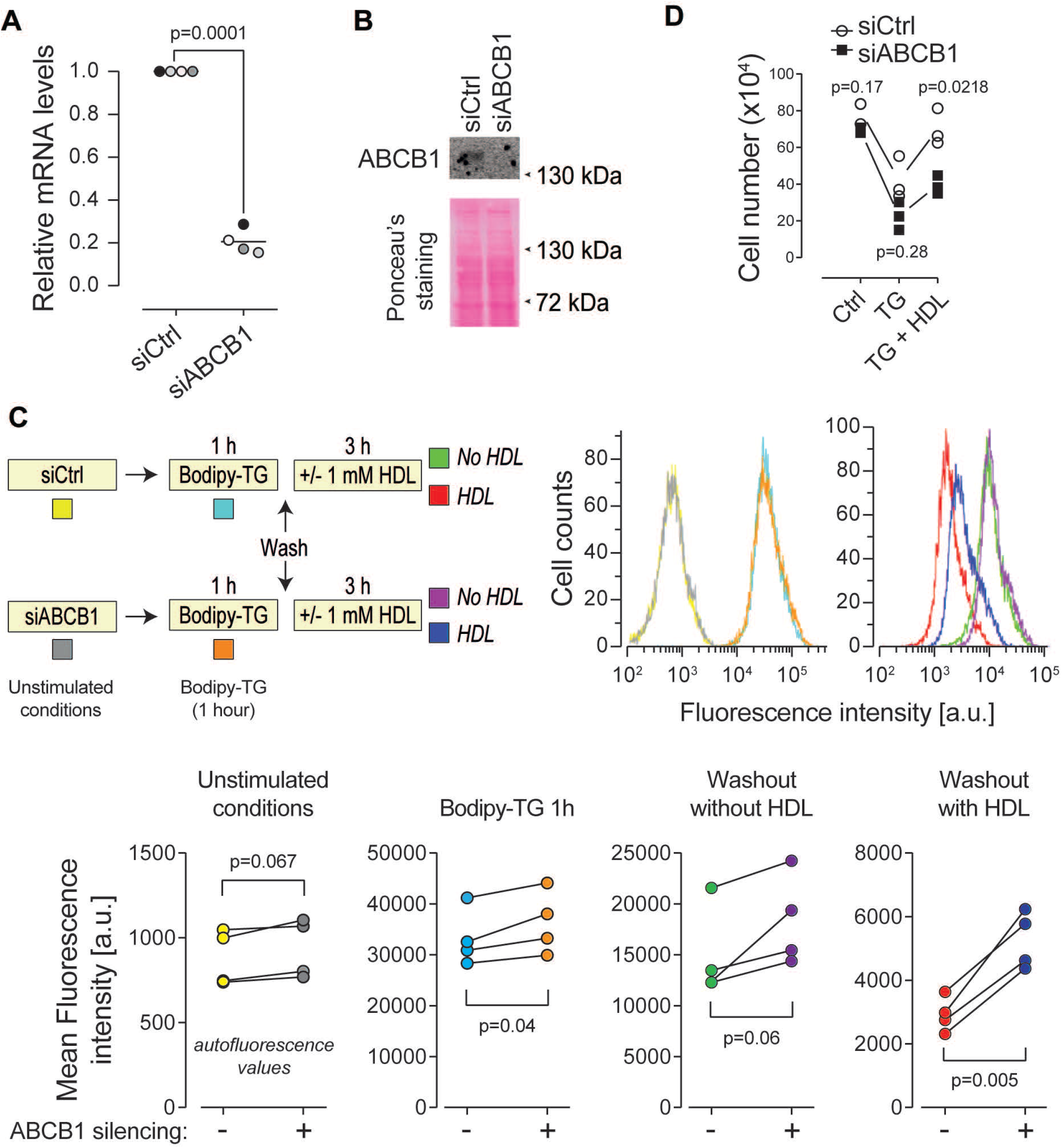
ABCB1 participates in HDL-mediated BODIPY-TG efflux. HEK293T cells (80’000 per well of 12-well plates) were cultured overnight and then transfected with a control siRNA pool (siCtrl) that does not target any human mRNAs and a siRNA pool directed at ABCB1 (siABCB1). The cells were analyzed 72 hours post-transfection. **A-B**. Knockdown efficiency at the RNA level was evaluated by RT-PCR (panel A) and at the protein level by western blotting (panel B). **C**. Control and ABCB1-silenced cells were incubated with BODIPY-TG and HDLs as indicated in the scheme. Cell-associated fluorescence was then assessed by flow cytometry. The quantitation of 4 independent experiments is depicted below the flow cytometry profiles. **D**. Alternatively, 48 hours post-transfection, the cells were treated with 20 μM TG for 24 hours in the absence or in the presence of 1 mM HDLs. Cells remaining attached to the plates (viable cells) were counted using light microscopy.

## Discussion

Low HDL levels are associated with the risk of developing several diseases including cardiovascular disease and diabetes [2, 28–30]. Whether high HDL levels directly participate in protective responses or whether they are mere markers of healthier metabolic states is a debated issue [31–33]. What is clear however is that HDLs can protect cells from a variety of toxic compounds [2]. Earlier work has suggested that HDLs activate intracellular pathways, such as those leading to Akt activation [34], to protect cells but this has recently been challenged [4]. The mechanisms allowing HDLs to protect cells are therefore still debated and require further characterization. The present work provides evidence that the drug efflux capacity of HDLs contributes to their protective effects against lipophilic drugs such as TG and staurosporine.

We previously showed that HDLs blocked the ER stress response induced by TG in pancreatic beta cells, efficiently protecting the cells from undergoing apoptosis [5]. HDLs were also shown to protect beta cells from tunicamycin, another ER stressor [6]. This protection occurred downstream of the ER stress response as HDLs in this case did not interfere with the capacity of tunicamycin to induce an ER stress response [6]. TG is lipophilic and, as shown in the present study, is readily extracted from cells by HDLs. HDLs therefore protect cells from TG by reducing the exposure of the cells to this drug. In contrast, tunicamycin is hydrophilic and is consequently not expected to be captured by HDLs. This can explain why HDLs do not prevent tunicamycin from activating an ER stress response. The way HDLs are inhibiting the death response induced by tunicamycin must therefore occur downstream of the activation of the ER stress response but how this occurs has eluded characterization so far.

HDLs are well-known cholesterol carrier. They can extract cholesterol from certain tissues (e.g. atherosclerotic plaques) but they can also deliver this lipid to other tissues, such as steroidogenic organs (e.g. adrenal glands, gonads). In some species, such as the mouse, HDLs are the main cholesterol carriers (in humans triglyceride-rich lipoproteins are the main cholesterol carriers) [35, 36]. The capacity of HDLs to take up or deliver cholesterol depends on how much cholesterol they already carry and the concentration of cholesterol found in membranes of the cells they interact with. By extension, whether HDLs give to or take from cells lipophilic drugs will depend on the respective levels of the drugs in the lipoprotein particles and in the plasma membrane. The capacity of HDLs to carry drugs has recently been discussed in the context of drug delivery [37]. HDLs can be viewed as natural nanoparticles that can be loaded with specific cargos bearing therapeutic properties [38]. Based on the finding reported in the present study, the drug efflux capacity of HDLs should be included in the design of strategies based on delivering therapeutic compounds to cells using HDL carriers. To illustrate this point, we can mention the observation that paclitaxel brought into cells by paclitaxel-loaded recombinant HDL particles is reduced by 70% by free HDL particles [9]. Consequently, when lipoproteins are utilized as drug carriers, drug dosage should be optimized to balance the effect of efflux to plasma HDLs.

The present work shows that HDLs are potent “extractor” of lipophilic compounds but are poor at promoting the efflux of hydrophilic drugs from cells (Figure S6). Hydrophilic drugs may therefore represent more ideal cargos to be transported by lipoproteins, as long as they can be loaded on lipoprotein particles [39].

HDLs can extract toxic xenobiotics such as TG, but can also promote the cellular efflux of therapeutic compounds such as the glibenclamide anti-diabetic drug (Figure 6). Determining the intrinsic capacity of HDLs from patients to capture a given drug may therefore inform clinicians on which drug to use, and at which dosage, for optimal therapeutic treatment.

Our data indicate that the ability to promote lipophilic drug efflux is not a unique specificity of HDLs, but is also present in two other serum components: albumin and LDLs. Albumin can enhance cellular cholesterol efflux by aqueous diffusion mechanism [40]. Albumin may employ similar mechanism to promote drug efflux, but we show here in the case of TG that this is far less efficacious than what can be achieved by lipoproteins (Figure S7). LDLs also contribute to ability of serum to promote TG efflux from cells but they are less potent than HDLs to do so (Figure S7). These results highlight the importance of controlling the impact of serum in experiments assessing the cellular activity of drugs and compounds, in particular those that are lipophilic.

Cholesterol efflux to HDLs is promoted by a series of ABC transporters. For example ABCA1 is involved in cholesterol transfer to nascent HDL particles and ABCG1 mediates the HDL cholesterol loading during the process of reverse cholesterol transport from arteries to the liver [41]. Our result indicate that ABCB1 participates in HDL-mediated lipophilic drug efflux. This efflux is however not solely dependent on ABCB1 as it still occurs with a delayed kinetics in the absence of this transporter. Other ABC transporters may share with ABCB1, at least partially, the capacity of promoting drug efflux to HDLs. However this is probably not a common feature of all ABC transporters as shown by the fact that ABCG2 silencing does not impact HDL-mediated TG cellular efflux (Figure S8). Another, non-exclusive possibility, is that there is a basal passive diffusion of drugs from the cell membrane to HDLs particles which can be enhanced by specific transporters such as ABCB1.

Clinical investigation is still needed to understand whether drug delivery to tissues and cells are causally associated or not with specific lipoprotein levels. Further research will have to be performed to evaluate *in vivo* how and to which extent HDLs extract hydrophobic compound and xenobiotics. This will expand our knowledge on the transport functions of lipoproteins beyond their classical physiological lipid biding capacity.

## Legends to Supplementary Figures

**Figure S1.**
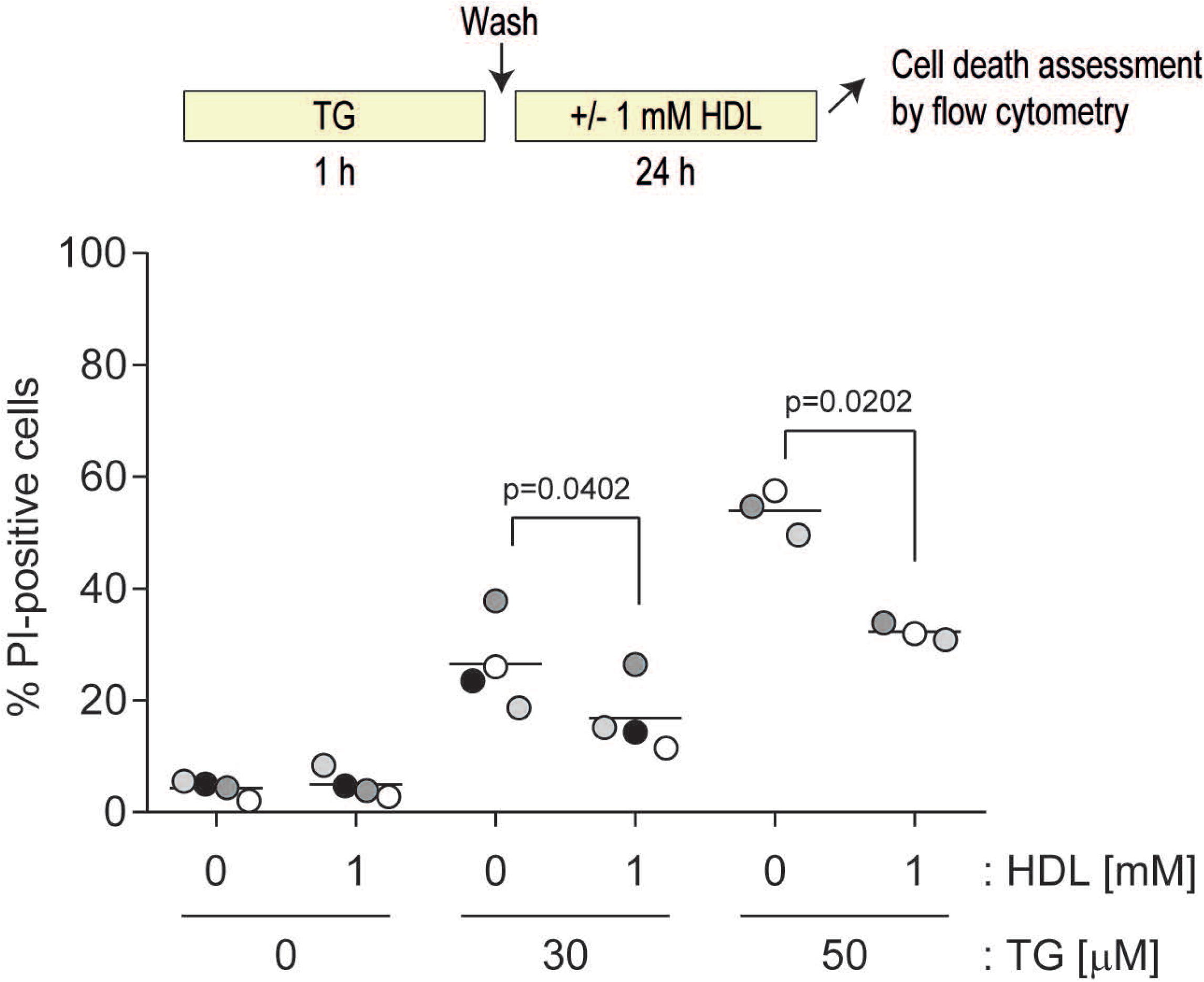
HDLs protect TG-exposed cells. DLD-1 cells were seeded in 6-well plates (100’000 per well). Twenty-four hours later, cells were pre-treated 1 hour with the indicated concentrations of thapsigargin (in μM). Cells were washed once with PBS and then incubated or not with 1 mM HDLs for an additional 24 hour period at which time cell death was assessed by PI incorporation using flow cytometry.

**Figure S2.**
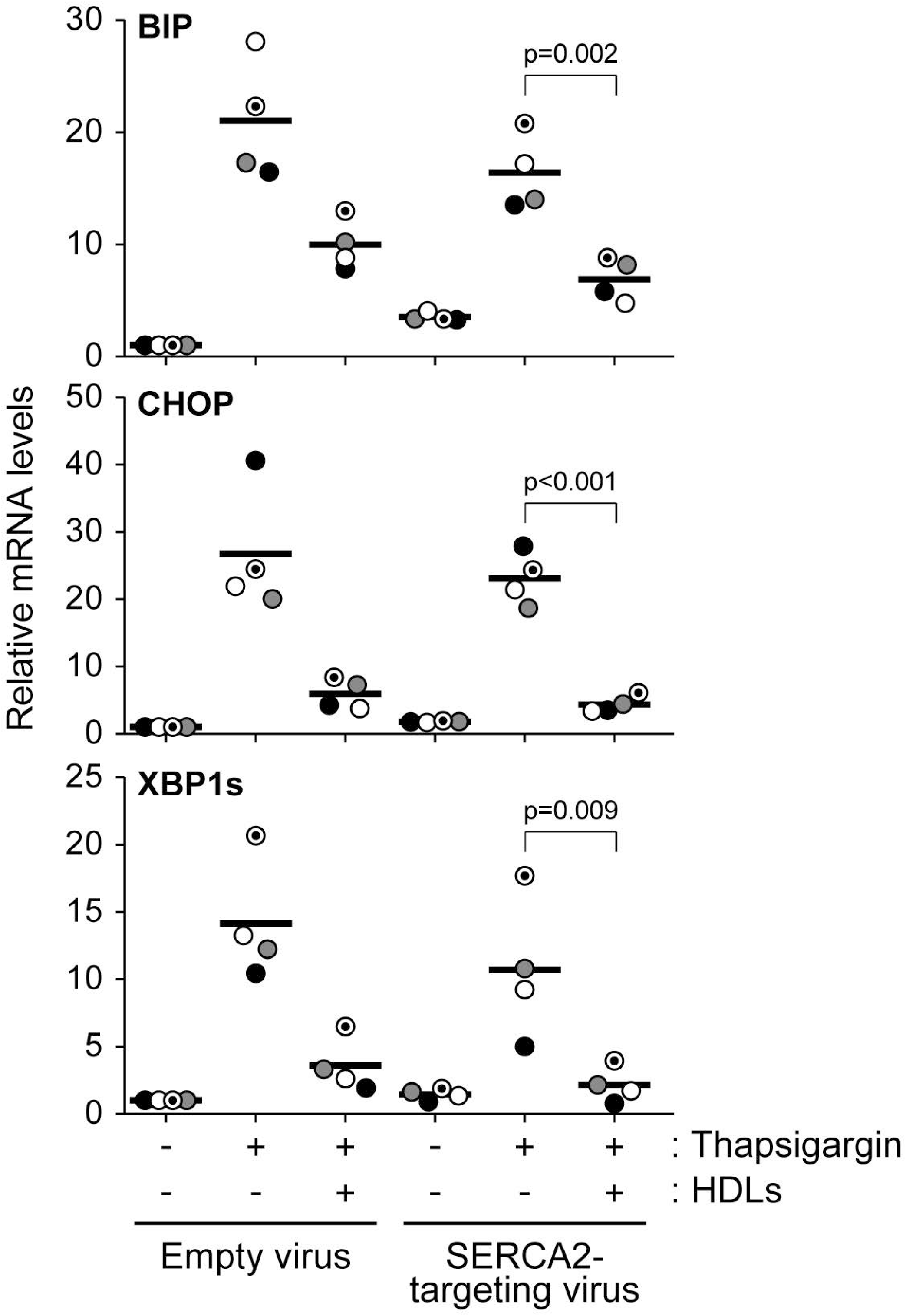
The effect of HDLs on thapsigargin-induced ER stress in SERCA2 knockdown DLD-1 cells. DLD-1 cells that were infected with a control virus or a lentivirus encoding for an shRNA directed against SERCA2 (see Figure 3B) were seeded in 6-well plates (200’000 per well). Twenty four hours later, cells were treated or not with thapsigargin (15 μM) in the presence or in the absence of 1 mM HDL for an additional 24 hour period. The mRNA expression levels of the indicated ER stress markers were measured by RT-PCR.

**Figure S3.**
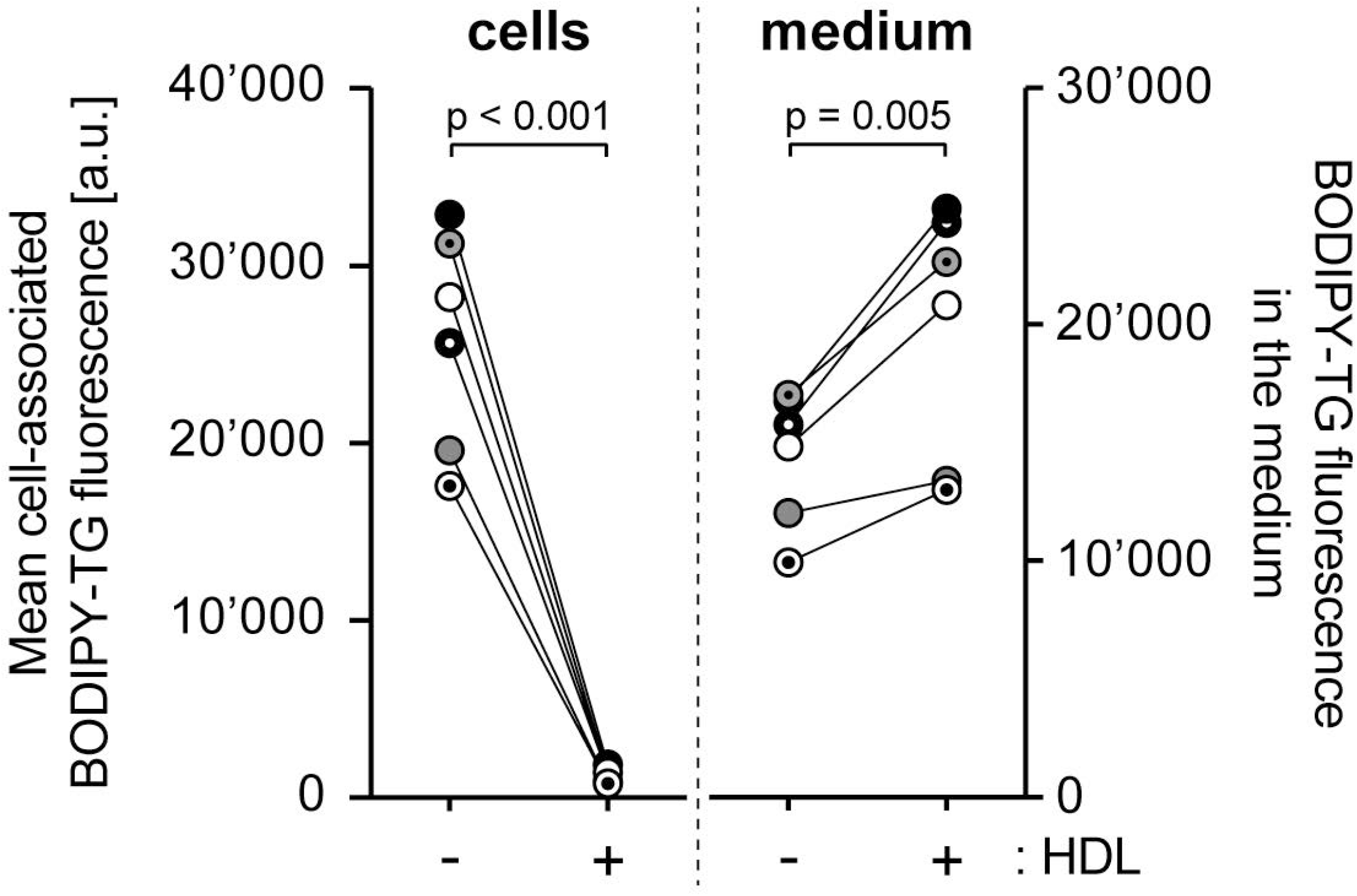
HDL-mediated BODIPY-thapsigargin extraction in HeLa cells. Hela cells (150’000 cells) were plated in 6-well plates. Twenty four hours later, they were treated with 1 μM BODIPY-TG for 1 hour, washed once with PBS, and then incubated in the absence or in the presence of 1 mM HDL for another 3 hours. The cell-associated BODIPY-TG levels were assessed by flow cytometry (left hand side graph labelled “cells”) and those in the medium measured using a Cytation 3 cell imaging multi-mode reader (excitation wavelength: 485 nm; emission wavelength: 538 nm) (right hand side graph labelled “medium”). The results are derived from 6 independent experiments (each labelled with a unique symbol).

**Figure S4.**
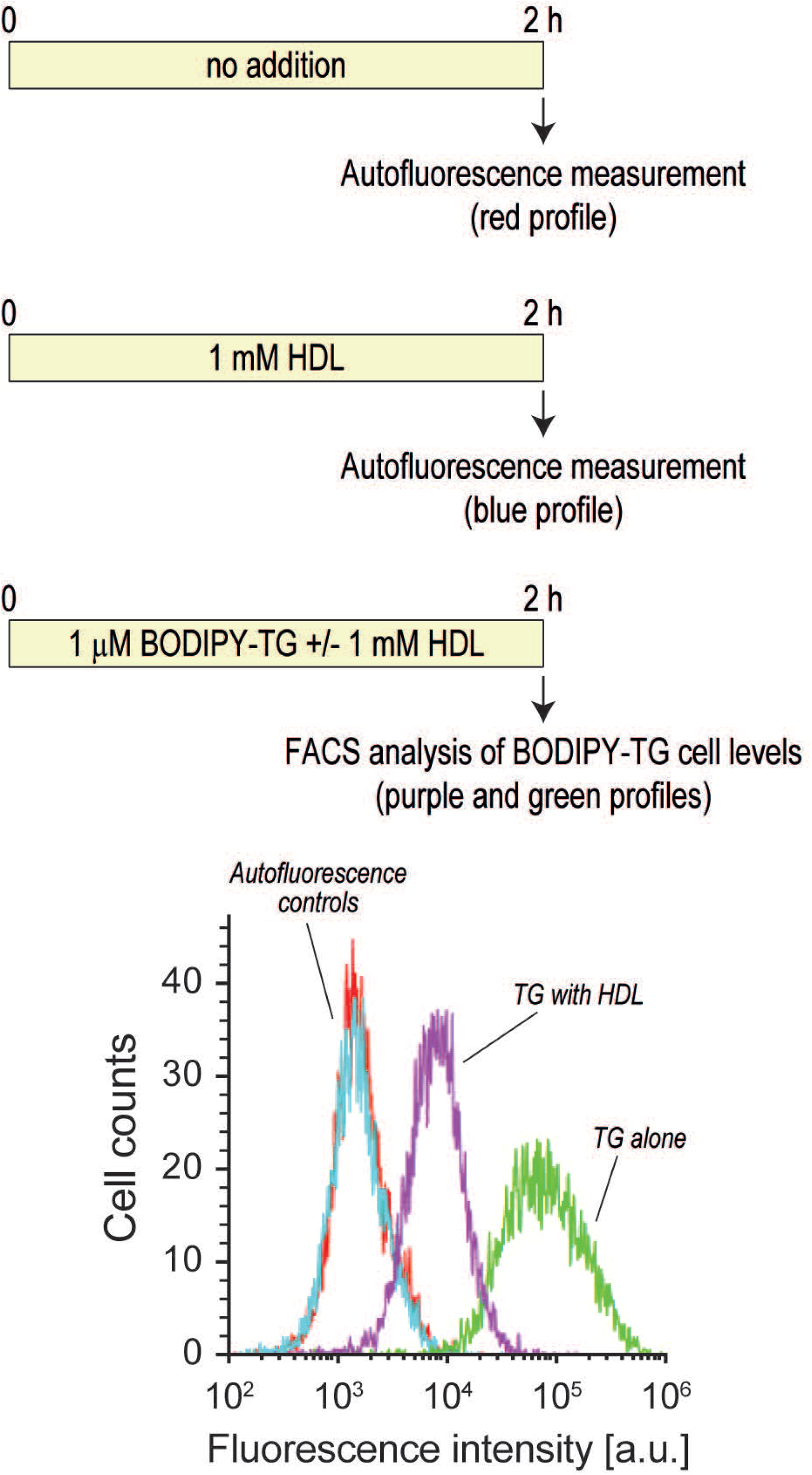
BODIPY-TG accumulation in cells in the continuous presence of HDLs. Min6 cells (300’000 cells per well) were treated with 1 μM BODIPY-TG in the presence or in the absence of 1 mM HDL for 2 hours. The cells were then collected and fluorescence intensity was monitored by flow cytometry. The autofluorescence profiles of untreated and HDL-incubated cells are also shown.

**Figure S5.**
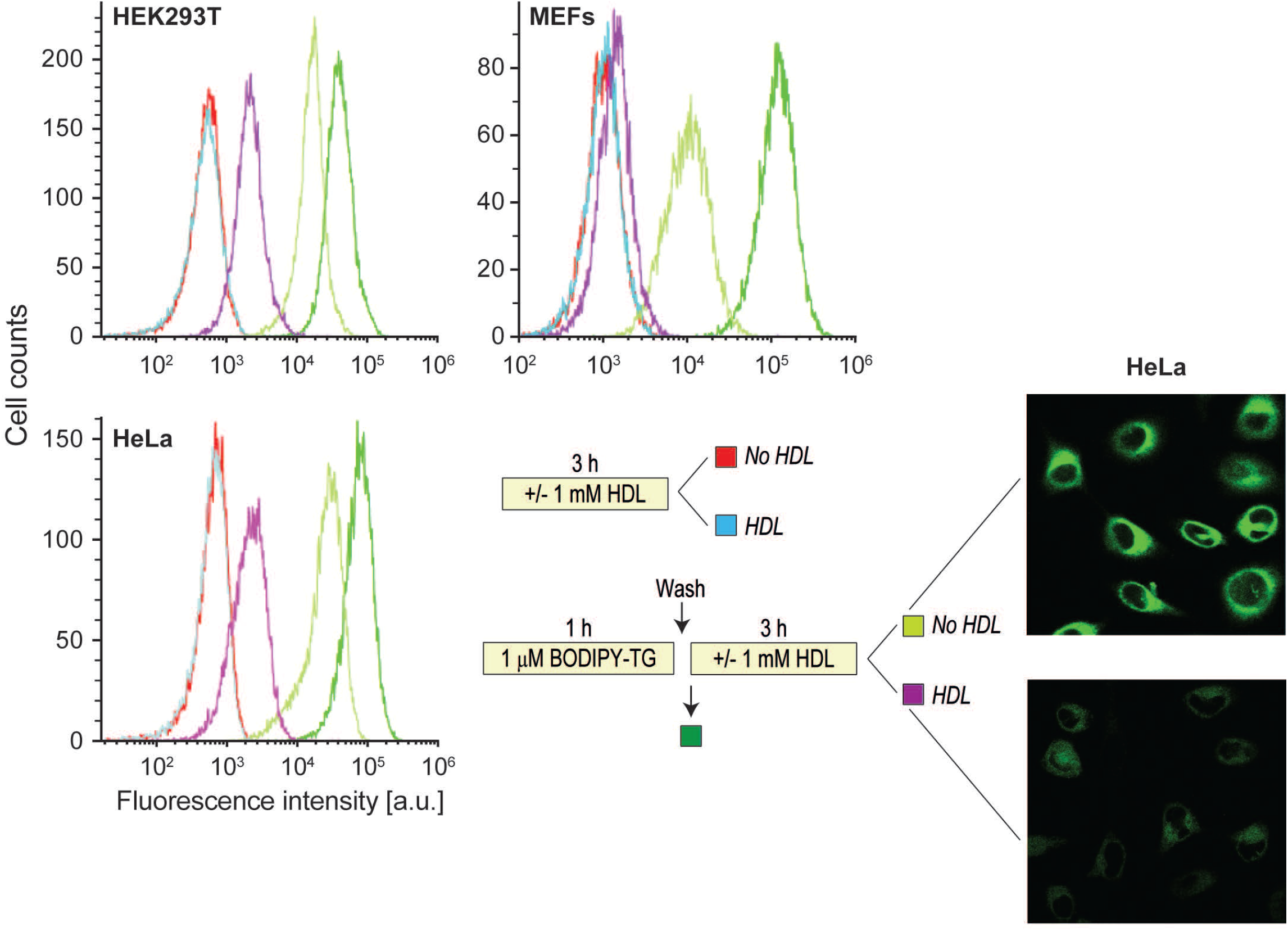
HDL-mediated BODIPY-TG efflux in various cell lines. Cells were seeded in 6 well plates (150’000, 80’000, and 200’000 per well for HEK293T, MEFs, and HeLa cells, respectively) one day before being treated as described in the scheme. Cells were then analyzed by flow cytometry. The images on the right hand side correspond to confocal imaging of 120’000 HeLa cells seeded the previous day in 35 mm glass bottom dishes and treated as shown in the scheme. The recording confocal settings between the two images were identical.

**Figure S6.**
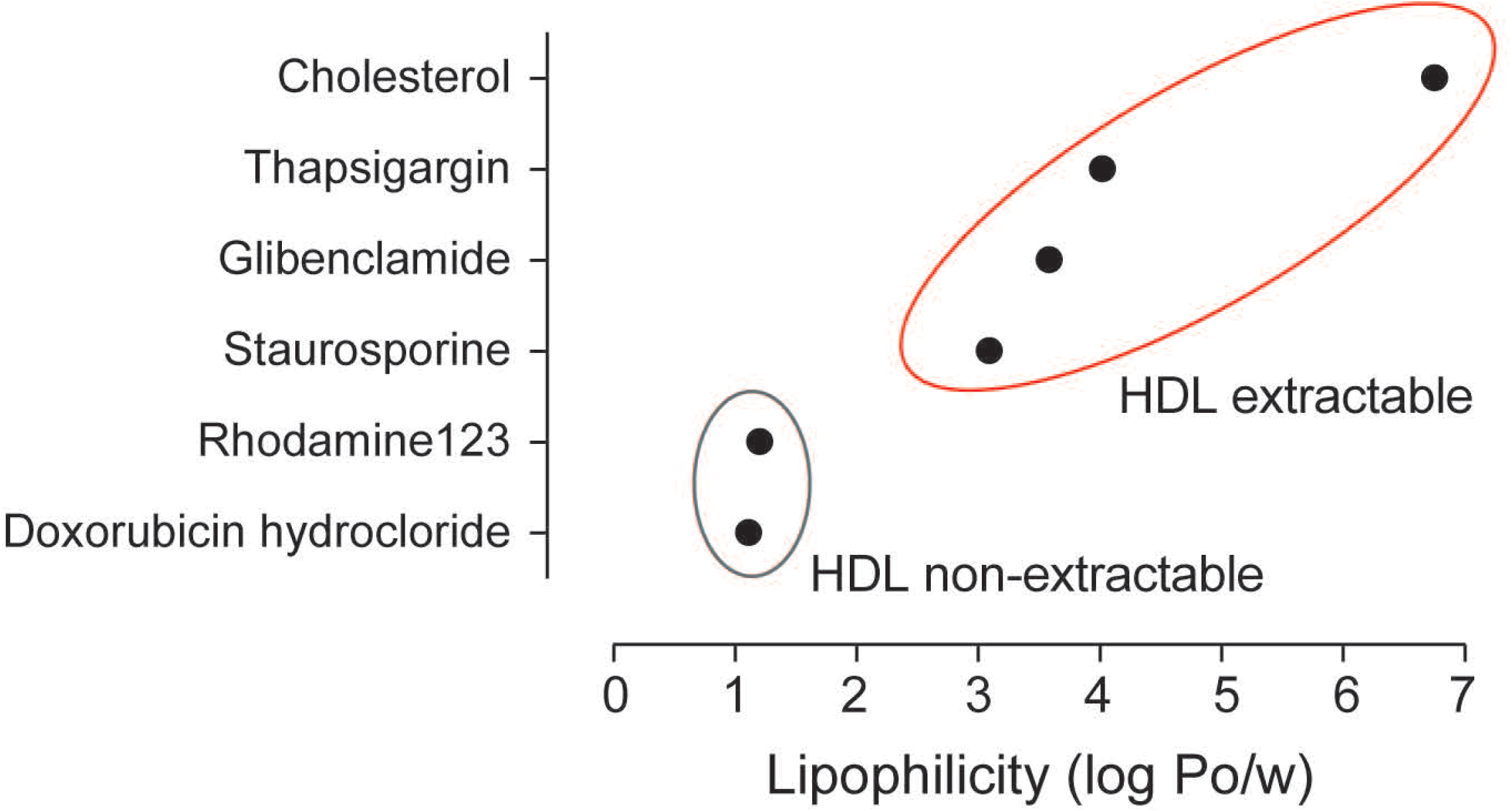
Correlation between HDL-mediated efflux efficiency and drug lipophilicity. The n-octanol/water partition coefficient (log P_o/w_) of the indicated compounds was obtained through the SwissADME website (http://www.swissadme.ch) [21].

**Figure S7.**
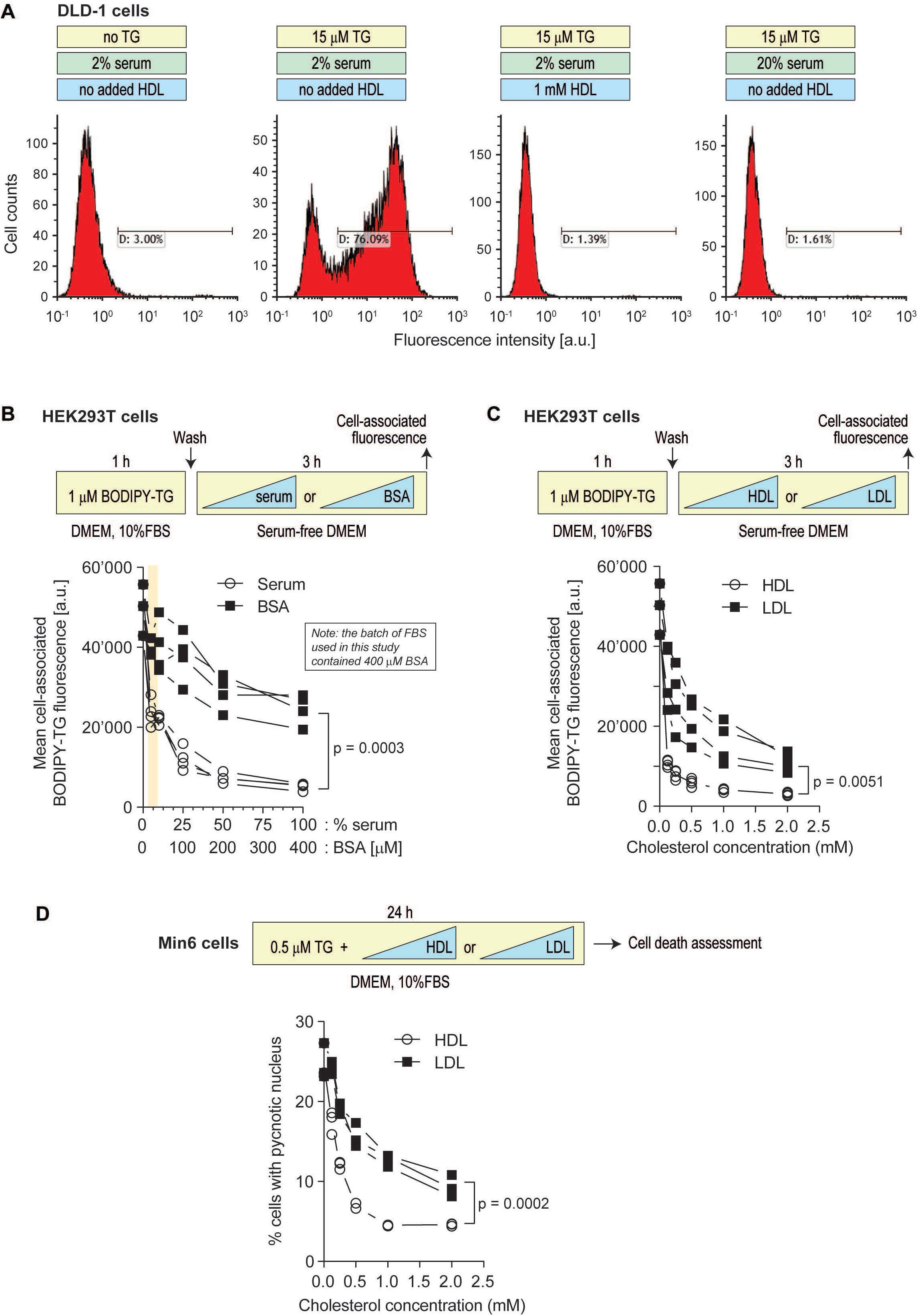
Serum components other than HDLs can extract TG from cells but less efficiently than HDLs. **A**. DLD-1 cells (100,000 cells per well) were treated as indicated in the scheme above the panel. Cell death was measured by PI staining. **B**. HEK293T cells (150,000 cells per well) were treated first 1 hour with 1 μM BODIPY-TG in DMEM, 10% FBS and then 3 hours with increasing concentrations of serum or BSA in serum-free DMEM. The concentration of BSA were chosen as to correspond to those found in serum (i.e. 0.4 mM BSA in 100% serum). Cell-associated BODIPY-TG was assessed by flow cytometry. **C**. Alternatively, instead of being incubated with serum or BSA, HEK293T cells were treated with increasing concentrations of HDLs or LDLs. The data are represented based on cholesterol content. **D**. Min6 cells (300,000 cells per well) were seeded in 6-well plates for 24 hours. Then cells were treated with 0.5 μM TG in the presence of increasing concentration of the indicated lipoproteins for 24 hours. Cell death was assessed by determining the percentage of cells with pycnotic nuclei.

**Figure S8.**
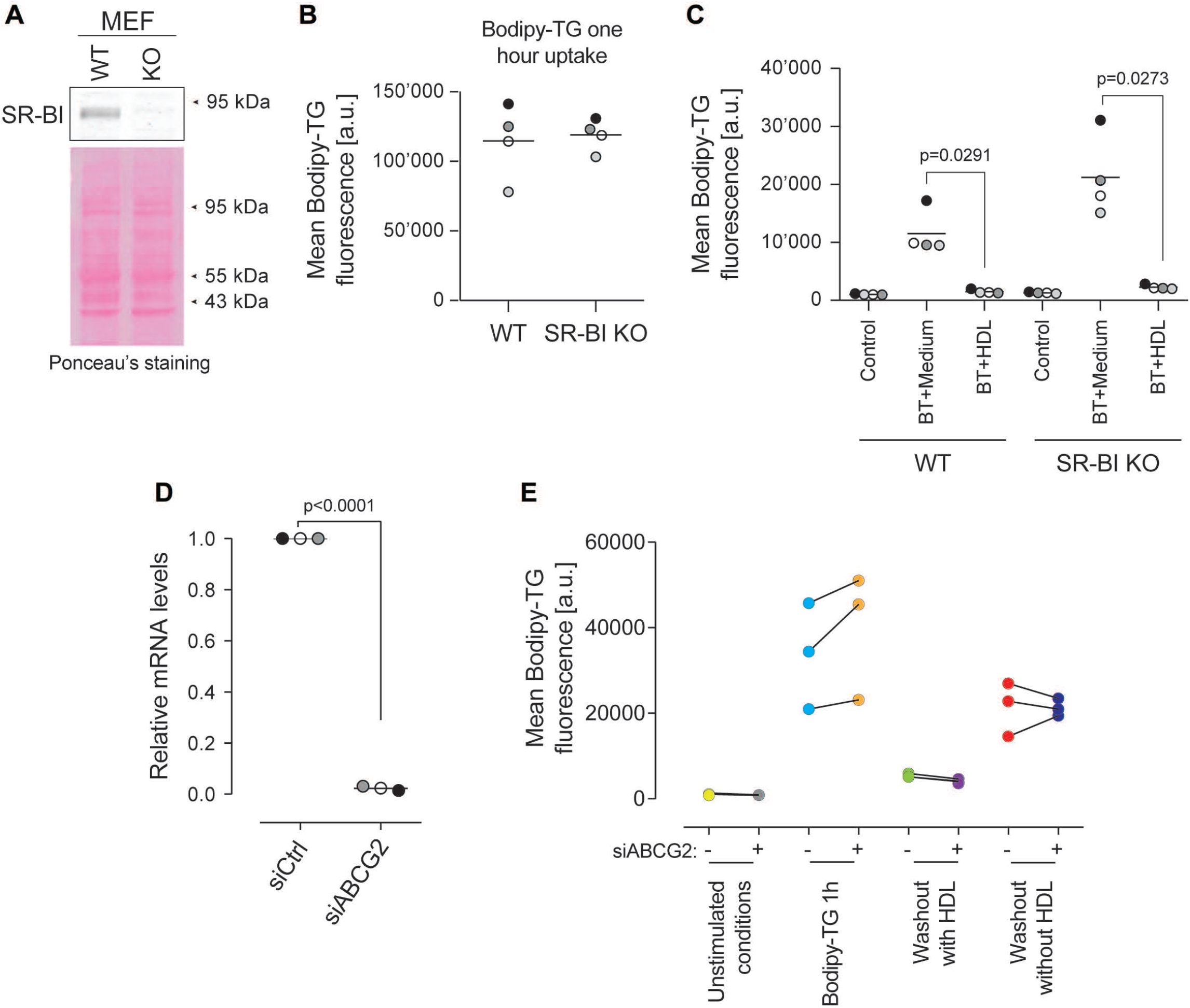
The effect of SR-BI and ABCG2 on BODIPY-TG efflux. **A**. SR-BI expression in wild-type (WT) and SR-BI knockout (KO) MEF cells. **B**. BODIPY-TG uptake in WT and SR-BI KO cells. **C**. BODIPY-TG efflux in WT and SR-BI KO cells. **D-E.** MCF7 cells (80,000 per well) were plated in a 12-well plate. Then cells were transfected with a pool of siRNA directed at ABCG2 and analyzed 72 hours post-transfection. Knockdown efficiency was assessed at the mRNA level by RT-PCR (panel D). Alternatively, cells were treated as indicated in the scheme shown in Figure 7C. Cell-associated drug fluorescence was measured by flow cytometry. The graphs present the data derived from 3 independent experiments (panel E).

## Acknowledgements

This work was supported by a Sinergia grant from the Swiss National Science Foundation (no. CRSII3_154420). We are thankful to the Cellular Imaging Facility platform at the University of Lausanne for the resources provided and their technical support.

